# Removing bias against short sequences enables northern blotting to better complement RNA-seq for the study of small RNAs

**DOI:** 10.1101/068031

**Authors:** Yun S. Choi, Lanelle O. Edwards, Aubrey DiBello, Antony M. Jose

## Abstract

Changes in small non-coding RNAs such as micro RNAs (miRNAs) can serve as indicators of disease and can be measured using next-generation sequencing of RNA (RNA-seq). Here, we highlight the need for approaches that complement RNA-seq, discover that northern blotting of small RNAs is biased against short sequences, and develop a protocol that removes this bias. We found that multiple small RNA-seq datasets from the worm *C. elegans* had shorter forms of miRNAs that appear to be degradation products that arose during the preparatory steps required for RNA-seq. When using northern blotting during these studies, we discovered that miRNA-length probes can have a ~360-fold bias against detecting even synthetic sequences that are 8 nt shorter. By using shorter probes and by performing hybridization and washes at low temperatures, we greatly reduced this bias to enable equivalent detection of 24 nt to 14 nt RNAs. Our protocol can better discriminate RNAs that differ by a single nucleotide and can detect specific miRNAs present in total RNA from *C. elegans*. This improved northern blotting is particularly useful to obtain a measure of small RNA integrity, analyze products of RNA processing or turnover, and analyze functional RNAs that are shorter than typical miRNAs.

## INTRODUCTION

Small non-coding RNAs such as microRNAs (miRNAs) regulate much of our genome (1) and thus can be broad indicators of health and disease. Next generation RNA sequencing (RNA-seq) (2) has quickly developed to become a powerful method to measure the expression of small RNAs. Because changes in miRNA expression have been observed in multiple disease states (3), there is great interest in measuring miRNA levels accurately for use in clinical diagnosis (4,5). Therefore, several additional approaches that augment RNA-seq for the accurate measurement of miRNA levels are being pursued (see ref. 6 for an overview).

A particular challenge in measuring miRNA abundance is that each miRNA gene can be expressed as a diverse population of related sequences (7) called isomiRs that differ at both 5’ and 3’ ends (8). Heterogeneity at the 3’ end was discovered early (9-11) and remains the dominant variation of miRNA sequence that is detected. These 3’ end variants can have both trimming and extension, which could be due to variation in miRNA precursor processing, activity of RNA exonucleases and/or non-templated nucleotide addition by nucleotidyltransferases (reviewed in 12). Modifications at the 3’ end (e.g. adenylation (13) and deadenylation (14)) have been associated with specific functional consequences. Recent pulse-chase experiments (15) have revealed that fast decaying miRNAs are associated with 3’ tailed and 3’ trimmed isomiRs. In contrast, the rarity of 5’ end variants has been attributed to the functional importance of the 5’ end seed sequence in determining the targets regulated by miRNAs (16). Yet, there are now reports that some isomiRs with different 5’ ends can behave as functionally distinct miRNAs and regulate different sets of target genes (17-20). Despite these advances, the biological significance of isomiRs, in most cases, remains unknown.

While miRNA variants are often first detected using RNA-seq, their validation and further study requires additional approaches to ensure that a detected RNA is present in the biological sample and not generated during the procedures required for RNA-seq. Although RNA-seq is sensitive, it has some limitations and sources of bias (reviewed in 21). For example, RNA-seq requires amplification, which can vary based on sequence composition (22,23), and ligation, which can vary based on modifications of terminal nucleotides (24) or presence of complementary sequences (25). In addition to these limitations that are intrinsic to RNA-seq, certain RNA purification methods can selectively miss sequences based on their composition (26). These considerations heighten the need for complementary approaches to accurately analyze populations of small RNAs.

One of the best approaches for validating the expression of a new RNA without the need for ligation or reverse transcription and irrespective of terminal modifications on the RNA is northern blotting. Therefore, substantial effort has been devoted to improving the sensitivity of this technique for the analysis of small RNAs. These include the use of locked nucleic acid (LNA)-modified probes (27), the use of chemical crosslinking of RNA to the membrane (28), and the demonstration that digoxigenin-labeled probes (29,30) can be more sensitive than radioactive probes (31). Together, improvements in sensitivity have made it possible to detect as little as 50 picomoles of a miRNA (31). Furthermore, the recent development of a technique to unmask small RNAs bound to complementary RNAs (25) and an approach to multiplex probes (32) continue to improve the ease of use and applicability of northern blotting.

Here we detect unexpected forms of well-studied *C. elegans* miRNAs in some RNA-seq datasets, discover that northern blotting can have drastic biases against short RNAs, and develop an improved northern blotting protocol that reduces this bias to enable detection of RNAs in the 14 nt to 24 nt size range.

## MATERIAL AND METHODS

### Analysis of RNA-seq datasets

All datasets analyzed in this study are listed in a Supplemental Table.

#### Detection of length variants of miRNAs in RNA-seq datasets

FASTQ or FASTA datasets were aligned using Bowtie to the *C. elegans* genome (Ce6) after removing adaptor sequences in the Galaxy interface (33-36). The bowtie parameters used were: bowtie -q -p 8 -S -n 2 -e 70 -l 28 – maxbts 125 -k 4 −phred33-<ce6 index> <fastq>. For adapter trimming, a minimum sequence length after trimming was set to 15 nt and all sequences with unknown (N) bases were discarded. Reads that aligned to genomic intervals of known miRNA genes (miRBase WBcel235) were filtered and used to calculate the average length of reads per miRNA gene. All reads (if total <1000) or 1000 randomly selected reads (if total >1000) that aligned to miRNAs that showed shortening were included to determine the coverage per nucleotide using Matlab (Mathworks).

#### Detection of miR-52 variants

Wild-type (N2) datasets that were deposited as compressed FASTA files or as SRA files were retrieved from the NCBI Sequence Read Archive. SRA datasets were converted to FASTQ format using SRA Toolkit V.2.4.2 and used regardless of quality scores. Prevalence of miR-52 with missing 5’ end nucleotides (5’-truncated) in N2 datasets was calculated by comparing the number of reads that exactly matched the full, annotated length (24 nt) to the number of reads that included the 3’-most 15 bases of miR-52 (ATATGTTTCCGTGCT) using *grep* function in unix. This 15-mer showed only one match to the *C. elegans* genome (miR-52) when searched with BLASTn. Only reads that exactly matched the reference genome were kept and reads with 3’ end variation were excluded. The number of reads required to consider a variant was scaled with the size of the RNA-seq dataset. In our detailed examination of mirR-52 variants, we counted only those forms of miR-52 that had a perfect match to the annotated 3’ end of miR-52. In addition to these 5’ truncated versions of miRNAs, there were additional forms that varied at both the 5’ and 3’ ends as expected because of the known 3’ end heterogeneity of miRNAs (12). Such miRNA variants with 3’ truncation and/or 3’ nucleotide addition were excluded from our counts.

Circularly permuted forms of miR-52 were identified by searching for reads (using *grep*) that exactly matched all the rearranged full-length miR-52 sequences. These sequences were used to search the nucleotide database with BLASTn and no alignments other than miR-52 were found in the *C. elegans* genome. All plots were created using Excel (Microsoft), Matlab (Mathworks) and Illustrator (Adobe).

### Generation of *mut-2(jam9)*

The *mut-2* locus (37) was edited by homology-based repair using the method described earlier (38,39). Briefly, plasmids that separately express Cas9, sgRNA, a homology repair template, and mCherry markers were co-injected into worms. Transgenic worms that expressed the mCherry marker were screened for edits in the *mut-2* locus.

The sgRNA targeting *mut-2* was designed using E-CRISP (40) and cloned into the pU6::unc-119 sgRNA expression vector (Addgene 46169, (39)). The *unc-119* sgRNA was removed from the plasmid by restriction digest (MfeI and XbaI; all restriction enzymes were from NEB). The large fragment of the plasmid digest was gel purified (28706, Qiagen) and used as the vector backbone for cloning of the new sgRNA. To prepare the insert containing the sgRNA targeting *mut-2*, two overlapping PCR products were amplified with Phusion DNA polymerase (NEB, used for all PCRs except as indicated) in separate reactions using pU6::unc-119 as a template. Primer pair YC001 and YC128 was used for the first reaction and primer pair YC127 and YC020 was used for the second reaction. Both PCR products were used as templates and the fused product was amplified using YC001 and YC020 as primers. The amplified product was column purified (28106, Qiagen), digested (MfeI and XbaI), gel purified (28706, Qiagen), and cloned into the prepared pU6::unc-119 sgRNA expression vector with T4 DNA ligase (NEB).

The homology repair template was prepared by initially cloning the *mut-2* locus and then replacing the *mut-2* coding regions with that of *gfp*. The *mut-2* coding sequence and ~1.4 kb of flanking sequences were amplified from wild-type (N2) genomic DNA using the primers YC087 and YC085. The PCR product was column purified (28106, Qiagen), digested (ApaI and PmlI), and used as the insert. The plasmid pCFJ-151 (Addgene 19330, (41)) was digested (ApaI and PmlI), and the ~2.3 kb fragment was gel purified, and used as vector. The resultant cloned plasmid (pmut-2_genomic) was used to amplify the 5’ (primers YC039 and YC134) and 3’ (primers YC137 and YC138) flanking sequences of *mut-2* in separate reactions. The *gfp* coding region was then amplified from pGC305 (Addgene 19646, (42)) with primers (YC135 and YC136) that also overlap *mut-2* flanking sequences. The three fragments were combined by overlap extension PCR using primers YC039 and YC138. The resultant ~1.8kb fusion product was gel purified, digested (PstI and BamHI), column purified (28106, Qiagen), and used as insert. The plasmid pmut-2_genomic was digested (PstI and BamHI), the ~3.8 kb fragment was gel purified (28706, Qiagen) and used as vector to generate pmut-2_to_eGFP_HRT.

Plasmids for microinjection were suspended in 10 mM Tris-Cl, pH8.5 to the following final concentrations: 50 ng/μl Peft-3::cas9-SV40_NLS::tbb-2 3’UTR (Addgene 46168, (39)); 22.5 ng/μl PU6::mut-2_sgRNA (based on Addgene 46169, (39)); 10 ng/μl pmut-2_to_eGFP_HRT; 10 ng/μl pMA122 (Addgene 34873, (43)); 10 ng/μl pGH8 (Addgene 19359, (41)); 2.5 ng/μl pCFJ90 (Addgene 19327, (41)); 5.0 ng/μl pCFJ104 (Addgene 19328, (41)).

F1 progeny of injected worms that expressed the mCherry coinjection markers were selected and allowed to have progeny. Twenty sets of five F2 progeny that did not express mCherry coinjection markers were moved to a plate. Pools of worms from each plate were screened for editing by PCR (primers YC176 and YC171). Individuals with successful editing were identified, homozygosed, confirmed by genomic DNA sequencing, and designated as *mut-2(jam9)*.

### RNA Purification

Worms grown on 35mm plates of LB agar and fed *E. coli* (OP50) were collected from starved plates by rinsing with M9 buffer (22 mM KH_2_PO_4_, 42 mM Na_2_HPO_4_, 86 mM NaCl, and 1 μM Mg MgSO_4_). Worms were pelleted (3 minutes at 750 × g tabletop microcentrifuge) and excess M9 buffer was aspirated leaving behind ~50 μl. 1.0 ml TRIzol (LifeTechnologies) was added to the worm pellet and frozen at -80°C. After thawing, the samples were processed according to manufacturer’s instructions. RNA was resuspended in nuclease free water (IDT) and its concentration was measured by UV spectrophotometry (Nanovue GE).

### RT-PCR

250 ng of total RNA was reverse transcribed with SuperScript III (LifeTechnologies) using 500 ng oligo-dT primer (IDT) according to manufacturer’s protocol. Gene specific primers and *Taq* polymerase were used to amplify target genes from 1 μl of the cDNA synthesis reaction. Cycling conditions were an initial incubation at 95°C for 30s followed by 30 cycles of: 95°C, 30s; annealing temperature (52°C, GFP; 55 °C, *mut-2*; and 55 °C, *ttb-2*), 30s; 72°C, 30s; and a final 72°C incubation of 5 min.

### Northern Blotting

Northern blotting was done as described earlier (31) with some essential modifications as noted below. In brief, 5 μg (*mut-2(jam9)*) or 10 μg (miRNA) of total worm RNA or 100 fmol of each synthetic RNA was separated by denaturing 20% PAGE (7M urea) and transferred (25 V, 1.0 A, 30 min, BioRad Transblot-Turbo) to a positively charged nylon membrane (11209299001, Roche). RNA was crosslinked to the membrane using UV irradiation (1200 μJ/m^2^, VWR UV Crosslinker). Synthetic DNA (IDT) probes were made with Roche DIG Oligonucleotide Tailing kit following manufacturer’s protocol or bought as 5’-monodigoxigenin-labeled (mono-DIG) oligonucleotides (IDT). Probes were hybridized to membranes (2.5 pmol probe in 5 ml ULTRAhyb buffer, Ambion LifeTechnologies) by incubating over night (with rotation at 8 RPM in a VWR Hybridization Oven) at 42°C for full-length probes and at room temperature (RT, ~22°C) for short probes. Membranes were then washed (2 × 15 min. at 37°C for full-length miR-52 probe; 2 × 15 min. at 42°C for full-length miR-53 probe; and 2 × 15 min. at RT for all short probes). Membranes were blocked (31), incubated with DIG antibodies (31), washed (4 × 15 min. at RT for long probes and 2 × 5 min. for short probes), developed (31), and imaged (Fujifilm LAS-3000). Blots were stripped by incubation in ~5-10 ml of 5% SDS at 80°C for 15 min. and washed twice with ~5-10 ml 2X SSC at 80°C for 5 min. The total RNA blot was probed first for miR-52, stripped, and then probed for U6. For the miR-52 & miR-53 blot, the probes were used in the following order: short miR-52, short miR-53, full-length miR-52, and full-length miR-53. The synthetic RNA blot was probed in the following order: miR-1, let-7, miR-58, and miR-52. The miRNA in total RNA preparations blot was probed in the following order: miR-52, miR-58, and U6. The blot for comparing mono-DIG labeling with DIG tailing was probed in the following order: 15 nt DIG-tailed, 15 nt mono-DIG, 24 nt DIG-tailed, 24 nt mono-DIG, and U6.

### Image analysis

Northern blot images were superimposed as separate layers in Photoshop (Adobe) and arranged so that the same area of the membrane was cropped for analysis. Cropped images were analyzed with ImageJ64 1.48v (44). Background was subtracted (50 pixels radius) before measuring band intensities using lanes of equal widths. Areas of peaks were determined by closing the base of each peak with a line segment. Peak areas were plotted with Excel and Illustrator.

### Worm strains

Wild-type (N2), *miR-52(4114)*, *miR-58(n4640)*, *mir-1(n4101)* and *mut-2(jam9[gfp])* were maintained at 20°C using standard methods (45).

## RESULTS

### Some miRNAs show substantial 5’ truncation in multiple RNA-seq datasets

A growing body of work shows that processes that change the length of a miRNA can be regulated (17-19) and are biologically important. To look for such variations in the length of miRNAs from *C. elegans*, we examined some published small RNA-seq datasets (Supplemental Table 1). Because non-templated nucleotide addition to 3’ ends of miRNAs is a common source of size heterogeneity (7), we compared datasets prepared from wild-type worms and from worms lacking a putative nucleotidyltransferase, MUT-2, which is predicted to add nucleotides to the 3’ ends of RNA (37). Surprisingly, we found that the average lengths of many miRNAs were substantially shorter in a dataset obtained from *mut-2(ne3364)* animals (Figure 1A), but to a lesser extent in a dataset obtained from *mut-2(ne298)* animals (data not shown). We also observed a similar shortening of miRNAs in a dataset from wild-type animals undergoing *pos-1* RNAi (Figure 1A) but not in a dataset from animals lacking MUT-16 (Figure 1A), which promotes formation of perinuclear foci where MUT-2 is thought to extend small RNAs (46,47). Despite the two datasets that showed an abundant population of shorter miRNAs being generated in different labs (see Supplemental Table 1), the datasets shared a subset of miRNAs that were substantially truncated. These included all members of an essential miRNA family in *C. elegans* – miR-51 to miR-56 (48) (Figure 1B for *mut-2(ne3364)* and wild-type animals undergoing RNAi). Furthermore, miRNA truncation was specific to the 5’ end of the more frequently sequenced arm (miRNA-3p or miRNA-5p) of the miRNA precursor (Figure 1C, Supplemental Figure 1).

**Figure 1.**
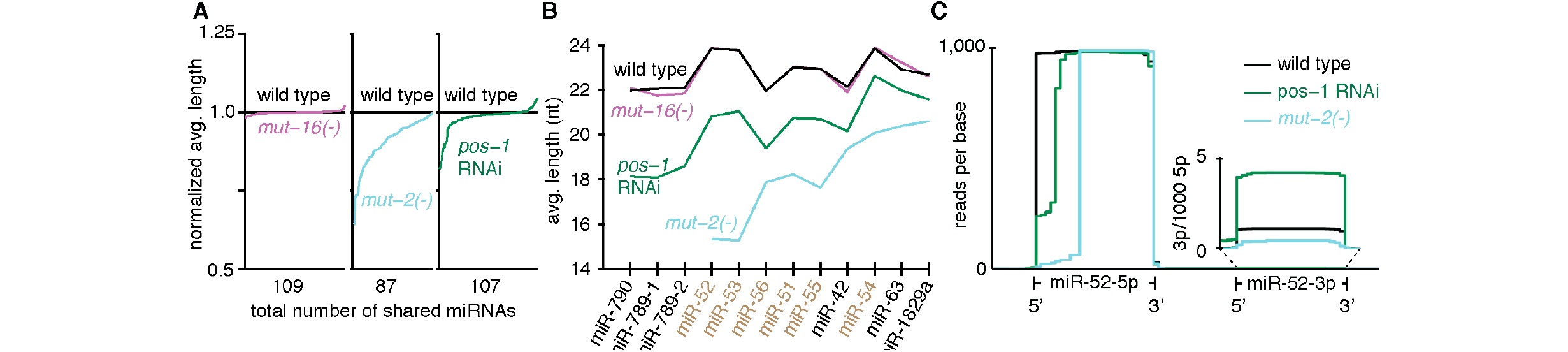
A subset of miRNAs is detected predominantly with missing nucleotides at the 5’ end in some small RNA-seq datasets. (**A**) Many miRNAs show altered average lengths in two different RNA-seq datasets. The average length of miRNAs detected in RNA-seq datasets both from wild-type animals (black) and from *mut-16(-)* (magenta), *mut-2(-)* (blue), or wild-type animals undergoing *pos-1* RNAi (green) were determined. The average length of each miRNA was normalized to that detected in the dataset from wild-type animals and plotted in rank order starting with the miRNAs that show the most proportional shortening. Total numbers of miRNAs present in both datasets (shared miRNAs) for each pair of datasets compared are indicated. (**B**) A common set of miRNAs, including the entire miR-51 to 56 family, show the most shortening in two different RNA-seq datasets. The average lengths of the most shortened miRNAs detected (A) were plotted for all datasets. Dataset labels as in (A) and the miR-51 to 56 family (brown) is highlighted. (**C**) The shorter length of miRNAs in both datasets is due to a shorter 5’ end and is specific to the more abundantly sequenced arm of the miRNA. 1000 RNA reads that map to a miRNA (miR-52 shown as an example) were sampled from wild-type (black), *mut-2(-)* (blue), or wild-type animals undergoing *pos-1* RNAi (green) and the resultant coverage (reads per base) of both arms of the miRNA gene (-5p and -3p, which are also in numbers per 1000 5p reads in insets) were plotted. Also see Supplemental Figure 1.

The 5’-truncated miRNAs could have been generated during the preparation of RNA for sequencing or during the lifecycle of miRNAs in vivo. The severely shortened forms observed in miRNAs from *mut-2(ne3364)* animals is surprising because *mut-2* has been reported to be required for silencing transposons through the generation of two other classes of small RNAs (22G and 21U) but not for the generation or stability of miRNAs (49). One possibility is that either this allele of *mut-2* or other mutations resulting from unregulated transposon activity are responsible for miRNA truncation.

### An abundant 5’-truncated miRNA observed in the *mut-2(ne3364)* RNA-seq dataset is not detectable by northern blotting in newly generated *mut-2(jam9)* null mutants

To test if the wild-type activity of MUT-2 is required to protect miR-51 to miR-56 family from 5’ truncation, we examined levels of the most highly expressed and most severely truncated member, miR-52, using northern blotting of total RNA prepared from *mut-2(-)* animals. To control for mutations induced by unregulated transposon activity in *mut-2(-)* animals (50), we analyzed three independent passages of a newly generated *mut-2* null mutant. Specifically, we replaced the entire coding portion of *mut-2* with DNA encoding *gfp* (Supplemental Figure 2A), which resulted in the expression of *gfp* instead of *mut-2* (Supplemental Figure 2B).

When miR-52 was examined in three independent passages of the *mut-2* null mutant using northern blotting, no evidence for shortening was detectable (Figure 2). For each passage, all detectable miR-52 – from wild-type and from *mut-2(-)* animals – was ~24 nt long, which is the annotated full length of miR-52. This result is in contrast to the 9 nt truncation that was observed in
~94% of the miR-52 reads by RNA-seq of *mut-2(ne3364)* worms (Figure 1C). Thus, our results suggest that MUT-2 is not required for the expression of full-length miR-52.

**Figure 2.**
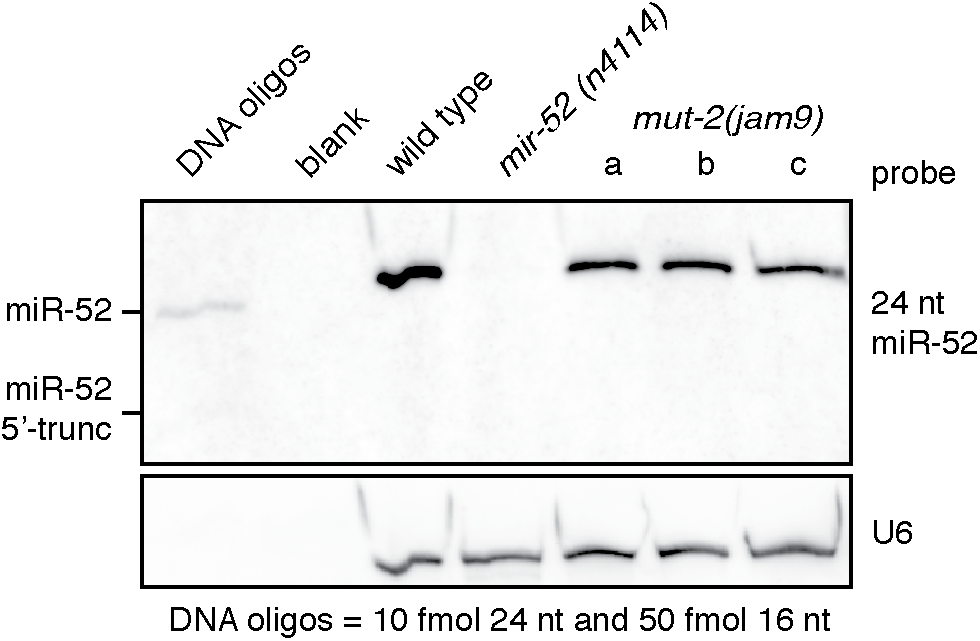
Newly generated *mut-2* null mutants do not show detectable shortening of miR-52 and short synthetic sequences are not effectively detected by conventional northern blotting. 5’ shortened versions of miR-52 are not detectable in *mut-2(jam9)* animals using conventional northern blotting. Total RNA prepared from wild-type, *mir-52* null mutant (*mir-52(n4114)*) or three isolates (a, b, c) of *mut-2* null mutant (*mut-2(jam9)*) animals was probed for miR-52 or U6 (loading control). When used as size standards (DNA oligos), 24 nt of mir-52 sequence (10 fmol) was detected more easily than 5X molar excess of 16 nt of 5’ shortened miR-52 sequence (50 fmol).

### 5’ truncated miRNAs are likely degradation products that arose during RNA-seq

Our failure to detect 5’-truncated miR-52 in *mut-2* null mutants raised the possibility that the apparent 5’ truncation observed in RNA-seq datasets results from the preparatory procedures required for RNA-seq. If so, then shorter versions of miRNAs may be detectable in additional RNA-seq datasets. Therefore, we looked for variants of miR-52 in 59 other datasets made from wild-type worms. These datasets were published by different labs that sequenced small RNA from worms of different stages or sexes using different protocols that did or did not select for a 5’-monophosphate for RNA sequencing. We observed 5’-truncated miR-52 in 17 of these 59 datasets with prevalence ranging from 79% to 19% of the miR-52 reads (Supplemental Figure 3A). Such 5’ truncation of miR-52 was not restricted to datasets from any lab, developmental stage, sex, or cloning method (Supplemental Figure 3A and Supplemental Table 1).

Not only was 5’-truncated miR-52 found in multiple RNA-seq datasets, all variations of circularly permuted miR-52 could also be detected (Supplemental Figure 3B). The simplest explanation for the presence of these circularly permuted sequences could be that the miRNA was circularized and then randomly cleaved along the phosphate backbone (Supplemental Figure 3C). If the linearized forms had the necessary 5’- and 3’-functional groups, it could then be captured in the RNA-seq library cloning protocol. Permuted miR-52 variants were rare compared to the total number of miR-52 reads but some forms were nevertheless sequenced more frequently than many other annotated miRNAs.

Taken together, these observations suggest that RNA-seq datasets contain 5’-truncated and other processed versions of miRNAs whose provenance is currently unclear.

### Northern blotting using miRNA-length probes is biased against detecting short RNAs

To complement RNA-seq data, additional methods that effectively detect small RNAs are needed. Although the average size of miRNAs is ~23 nt, the recent release of miRbase (V.21) contains a wide range of miRNA sizes (from the 15 nt miR-7238-3p from *Mus musculus* to the 34 nt miR-5971 from *Haemonchus contortus*) with varying degrees of functional validation. Therefore, a method that can effectively detect RNAs in the range of 15 nt to 34 nt would be useful to independently evaluate the expression of some miRNAs and other short RNAs like the truncated variants that we detected in RNA-seq datasets.

When examining the levels of the 24 nt long miR-52, we made an observation that suggests that typical northern blotting has a strong bias against shorter sequences. The 24 nt probe used was able to detect 10 fmol of 24 nt DNA analogue of miR-52 effectively but barely detected 50 fmol of 16 nt DNA representing 5’-truncated miR-52 (Figure 2, DNA oligos lane). Because both these DNA sequences were synthetic, the difference in their detection is likely due to differences in the length of the sequence and not due to any chemical changes to the nucleotides that could have occurred in vivo. Thus, northern blotting needs to be modified for effectively detecting short RNAs.

### Northern blotting can be improved to reduce bias against short RNAs and maintain target specificity

A previous study reported the detection of a specific ~14 nt RNA called pRNA in samples prepared from two species of bacteria using a modified northern blotting protocol (51). This approach used Locked Nucleic Acid (LNA) probes that are 5’digoxigenin-labeled, chemical crosslinking of RNA to nylon membranes using 1-ethyl-3(3-dimethylaminopropyl)-carbodiimide (EDC), native PAGE, and hybridization at 50°C to 68°C. While this approach is powerful, the authors noted a few shortcomings: (1) LNA probes are expensive and detection is affected by the distribution of LNA modifications on the probe; (2) EDC linking requires a terminal phosphate, in the absence of which, the signal was reduced by ~10 fold; and (3) to avoid EDC crosslinking to urea, RNA samples have to be run on native PAGE, which can affect the migration of small RNAs with secondary structure or that interact with complementary sequences (e.g. stable 6S RNA:pRNA hybrids). Therefore, we wondered whether the detection of miRNAs as well as truncated forms of miRNAs or other such shorter RNAs using cheaper DNA probes and under denaturing conditions could be achieved by performing hybridization at lower temperatures with UV crosslinking.

To begin testing this possibility, we used two different length probes to detect progressively 5’-shortened versions of synthetic miR-52 and miR-53. In addition to the 24 nt probes, we designed shorter probes (15 nt for miR-52 and 16 nt for miR-53) that target the 3’ end of both miRNAs and include the single nucleotide polymorphism (SNP) that distinguishes miR-52 from miR-53 (Figure 3A). To account for the differential base pairing potential of different length probes, we hybridized at 42°C for 24 nt probes and at room temperature (~22°C) for short probes. The 24 nt probe could only detect the 24 nt to 18 nt versions of the miRNAs, with rapidly diminishing signal intensity for shorter RNAs (Figure 3A & B, black). However, the short probes were able to detect all of the different length RNAs with comparable efficiency (Figure 3A & B, red). The stark bias against shorter RNAs when 24 nt probes were used (e.g. ~100-fold for 18 nt miR-53 and ~360-fold for 16 nt miR-52) was greatly reduced when 16 or 15 nt probes were used (a maximum of ~4-fold difference across 24 nt to 14 nt).

Another important consideration for northern blotting is specific recognition of a target RNA. We measured the specificity of short and long probes by comparing their ability to detect perfectly complementary RNA (target) with that to detect RNA that differed by a single nucleotide. The 24 nt miR-52 probe showed almost equal reactivity to 24 nt miR-52 and miR-53 (Figure 3A), consistent with previous reports of cross-reactivity (52). The 24 nt miR-53 probe, which was subjected to washes at higher temperature (42°C washes compared to 37°C used for miR-52), showed reduced cross-reactivity. However, the increased stringency of the washes increased the bias against detecting shorter RNAs as evidenced by the exponential decrease in signal as the length of target RNA decreased (Figure 3A & B). In contrast, short probes were able to detect all targets with comparable or better specificity than the full-length probes (Figure 3A & B and Supplemental Figure 4). Finally, short probes were able to specifically detect different small RNAs of varying GC content (Figure 3C). A 15 nt probe with only 27% GC could still bind specifically to its miR-1 target with no cross-reactivity to three other abundant synthetic RNAs on the same membrane (Figure 3C).

**Figure 3.**
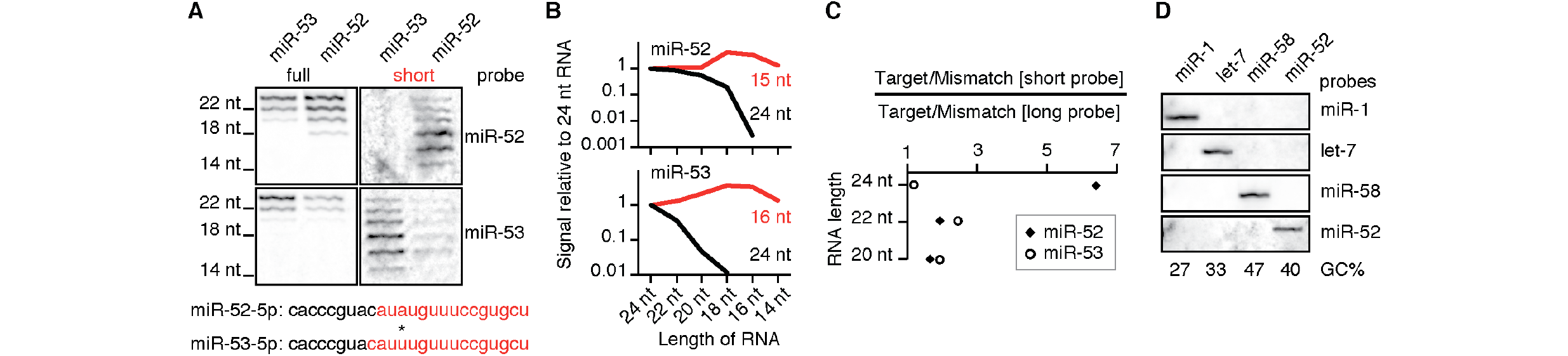
Short probes enable effective detection of short RNAs using northern blotting. (**A** and **B**) Short probes reduce length bias in northern blotting and enable detection of RNAs as short as 14 nt with single nucleotide discrimination. Equimolar (100 fmol) amounts of synthetic RNA representing miR-52 and miR-53 (differs from miR-52 by a single nucleotide, asterisk) with progressive 5’ shortening in 2 nt increments were analyzed by northern blotting using short probes (red, 15 nt or 16 nt) or using full-length probes (black, 24 nt). Northern blot (A) and its quantification (B) are shown. Relative intensity of each band was determined using that of 24 nt RNAs as reference (long and short probes are as in (A)). While blotting using full-length probes showed an extreme bias for longer RNA and substantial cross-reactivity (also see ref. 34), blotting with short probes detected all sizes with less cross-reactivity. (**C**) The selectivity of short DNA probes is as good or better than that of long DNA probes. The selectivity of a probe was calculated as the ratio of the signal from precisely complementary RNA (target RNA) to that from 1 nt mismatched complementary RNA (mismatch RNA) of the same length. The relative selectivity of the short probe (x-axis) compared to the long probe was calculated by taking the ratio of the ratios. (**D**) Short probes with differing GC content are able to selectively detect different miRNAs. One membrane with equimolar (100 fmol) amounts of synthetic RNA representing miR-1, let-7, miR-58, and miR-52 was sequentially probed using the corresponding
15 nt probes with the indicated GC content (% GC).

Thus, short probes and hybridization at low temperatures can reduce the bias against the detection of short RNAs that arise when long probes and typical hybridization temperatures are used.

### 15 nt probes can specifically detect miRNAs in total RNA from *C. elegans*

Although our method demonstrates the specific detection of synthetic RNAs, it will be of limited utility unless it can be applied to detect specific sequences in more complex mixtures of RNA. Therefore, we attempted the detection of three specific miRNAs (miR-1, miR-42, and miR-58) with 15 nt probes of varying GC content in preparations of *C. elegans* total RNA. These miRNAs were chosen based on the availability of viable deletion mutants that can be used as essential controls and to cover a range of GC content (miR-1, 27% GC; miR-52, 40% GC, and miR-58, 47% GC). Total RNA from wild-type worms and from mutant worms with a deletion for a miRNA gene was purified and used for northern blotting. Both miR-58 and miR-52 were detected in total RNA from wild-type worms but not in total RNA from the corresponding mutants (Figure 4). However, miR-1 could not be detected in any case despite seven attempts (data not shown), possibly because the low GC content of the probe
used (27% GC) is below the threshold for stable binding at room temperature. These results demonstrate the utility of the modifications we propose – shorter probes and lower hybridization temperatures – for the effective detection of a specific small RNA in total RNA preparations from an animal.

## DISCUSSION

We detected unexpected forms of miRNAs in some RNA-seq datasets and found that northern blotting using miRNA-length probes can be biased against detecting shorter RNAs. We demonstrate that adjusting probe size and hybridization conditions can allow equivalent detection of
24 nt to 14 nt RNAs with single-nucleotide sequence discrimination using affordable DNA oligos as probes.

### Possible origins of miRNA variants in RNA-seq datasets

We detect two unusual miRNA variants in RNA-seq datasets - 5’ truncated forms of miRNA and circularly permuted forms of miRNA.

The truncated forms of miRNA could arise during sample preparation before RNA extraction. We observed that in most cases truncated miRNAs were missing sequences from the 5’ end (Figure 1 and Supplemental Figure 1). Such truncations were particularly pronounced in all members of the miR-51 family. These observations suggest possible degradation of miRNAs by a 5’ to 3’ exoribonuclease. Consistently, *C. elegans* has a 5’ to 3’ exoribonuclease called XRN-2 that can act on miRNAs in general (53) and for which the miR-51 family could be an especially good substrate (54). Such mechanisms that can degrade miRNAs raise concerns about the accuracy of inferring the levels of miRNAs that function *in vivo* from RNA-seq and suggest the use of truncated versions of miR-51 family members as indicators of poor RNA quality.

Circularly permuted forms of miRNA could arise during ligation steps required for library preparation. The number of reads that we count as circularly permuted is likely an underestimation. The ~0.16% of miR-52 detected as circularly permuted in Supplemental Figure 3B reflect only the sub-population of circularized RNAs that became linear and were then successfully joined to sequencing adaptors. Those RNAs that remained as circles or did not get ligated to adaptors would escape detection. Although circular RNAs that result from RNA splicing have been detected in *C. elegans*(55,56), there is no evidence yet for the generation of circular miRNAs in vivo.

### Technical considerations for the use of northern blotting to complement RNA-seq

Our results suggest that the hybridization temperature and GC content of probes are critical variables for the detection of short RNAs. Long probes hybridized at 22°C had off-target binding and retained a bias against short RNAs (Supplemental Figure 4), while short probes hybridized at 37°C - 42°C did not have any detectable signal (data not shown). We also noted that hybridization below 20°C results in components of the hybridization buffer becoming insoluble. We failed to detect miR-1 using 15 nt probes with 27% GC, suggesting a requirement for a minimum number of GC base-pairs for stable binding with complementary RNA. Indeed, the miR-52 probe with 40% GC had a noticeably weaker signal compared to the miR-58 probe with 47% GC (Figure 4) – a difference of one GC base pair.

**Figure 4.**
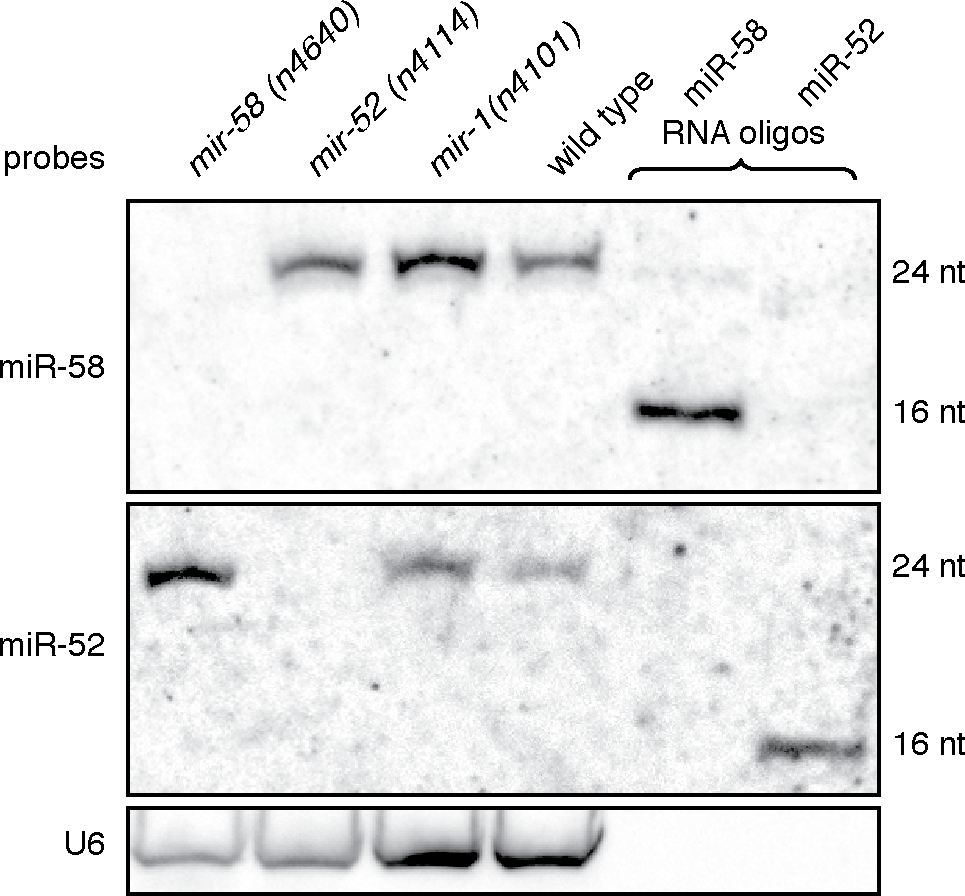
Short probes can specifically detect miRNAs in total RNA. Total RNA prepared from wild-type and miRNA deletion (*mir-58(n4640)*, *mir-52(n4114)*, and *mir-1(n4101)*) worm strains was used for northern blotting with short (15 nt) probes against mir-58 or mir-52. Synthetic RNAs that matched the sequence of full length (10 fmol of 24 nt, weakly detected) and shortened forms (50 fmol of 16 nt, clearly detected) of miR-58 and miR-52 were run as size markers. U6 was probed to indicate loading.

We found that low incubation temperatures and appropriate washing conditions can enable short sequences to specifically detect complementary RNA. This specificity was despite the addition of a long tail (>100 nt) of random dA and DIG-conjugated dU to generate the DIG-tailed probe. Mono-DIG-labeled 15 nt sequence probes failed to detect miR-52 and mono-DIG-labeled 24 nt sequence probes detected miR-52 with a weaker signal than DIG-tailed probes (Supplemental Figure 4). Thus, using DIG-tailed probes reduces cost (~20× compared with mono-DIG probes) and enhances the detectable signal without apparent off-target interactions from the long tail.

### Applications of the modified northern blotting described in this study

The drastic bias against short RNAs observed when miRNA-length probes are used (Figure 3A), suggest that the careful study of isomiRs (12) will require designing probes that match the shortest form of the miRNA. Furthermore, this bias raises the possibility that RNA quantified by northern blotting in previous studies might have under-reported RNAs shorter than the length of the probe. Such shorter RNAs could include biologically relevant RNAs such as product RNAs (~14 nt) in bacteria (57), NU RNAs (<18 nt) in *C. elegans* (58), or unusually small RNAs (~17 nt) that were reported to associate with Argonaute proteins in mammalian cells (59). The ability to better discriminate RNAs that differ by single nucleotides when using short probes (Figure 3C) could help resolve expression from two different alleles without the amplification necessary for RNA-seq (60) or PCR (61) approaches. Finally, reducing the bias against shorter RNAs could enable tracking intermediates of RNA processing or small RNA turnover (62) in mechanistic studies and is thus an essential consideration when probing the realm of tiny RNAs.

## ACKNOWLEDGEMENT

We thank members of the Jose lab for critical reading of the manuscript. We thank the *C. elegans* Genetic stock Center for worm strains. This manuscript has been improved with comments from anonymous reviewers who helped place this work in the context of other efforts underway to improve the detection of small RNAs.

---

**Supplemental Figure 1.**
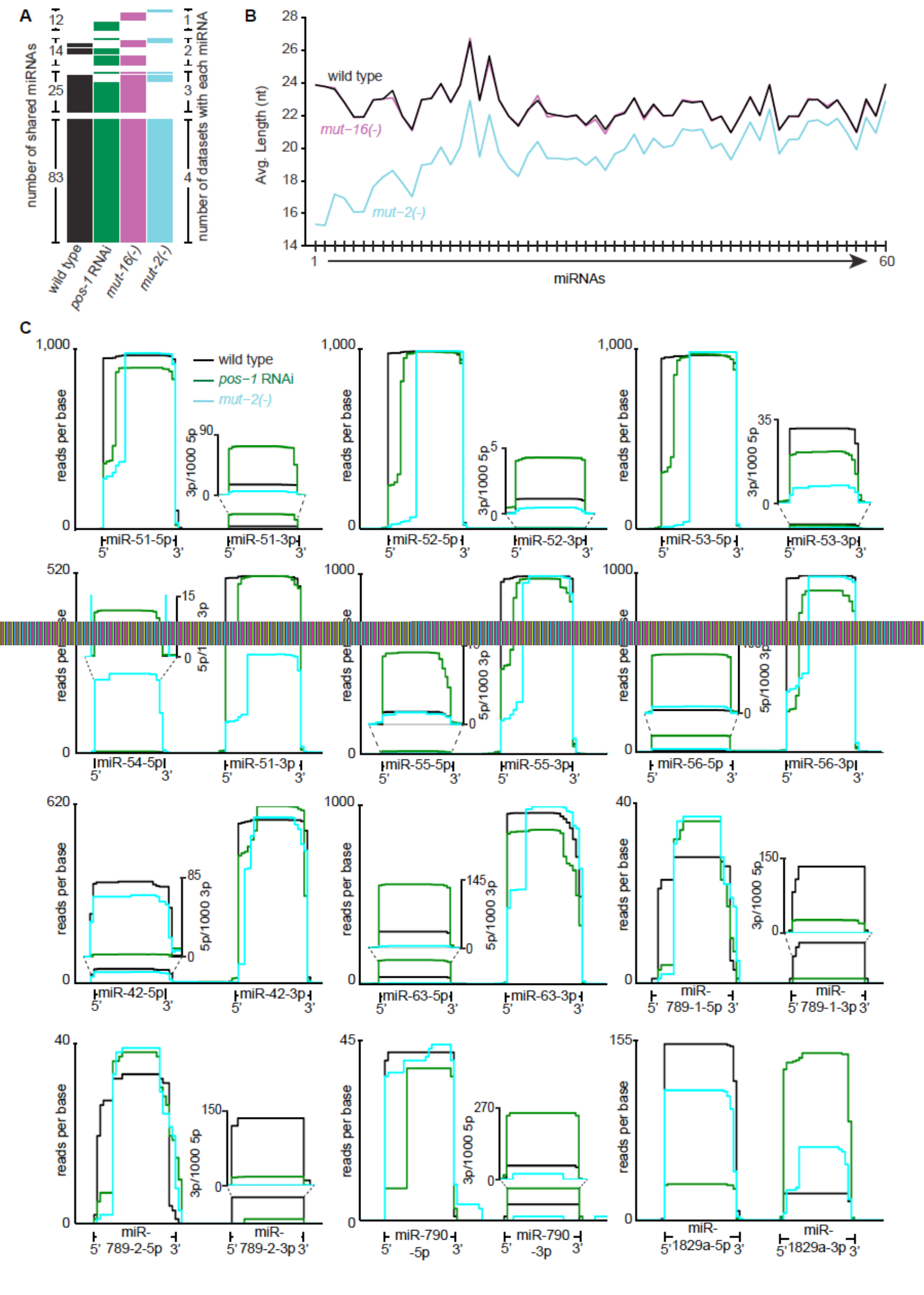
Some miRNAs are detected as shorter versions predominantly missing 5’ end nucleotides in two different small RNA-seq datasets. (**A**) 83 miRNAs are detected in four different datasets and thus their characteristics can be compared across datasets. Number of miRNAs present in one, two, three, or four RNA-seq datasets when datasets from wild-type animals (black), *mut-16(-)* animals (magenta), *mut-2(-)* animals (blue), or wild-type animals undergoing *pos-1* RNAi (green) were compared are shown. (**B**) Many miRNAs show altered average lengths in a *mut-2(-)* RNA-seq dataset. The average lengths (nt) of the miRNAs that were shortened by at least 1 nt on average in a *mut-2(-)* dataset were rank ordered and plotted from wild-type (black), *mut-16(-)* (magenta), and *mut-2(-)* (blue) datasets. (**C**) The shorter length of the twelve most shortened miRNAs in the *mut-2(-)* dataset appear to be due to missing 5’ end nts and to be specific to one of the sequenced arms of the miRNA. The plot for each miRNA is as described in Figure 1C.

**Supplemental Figure 2.**
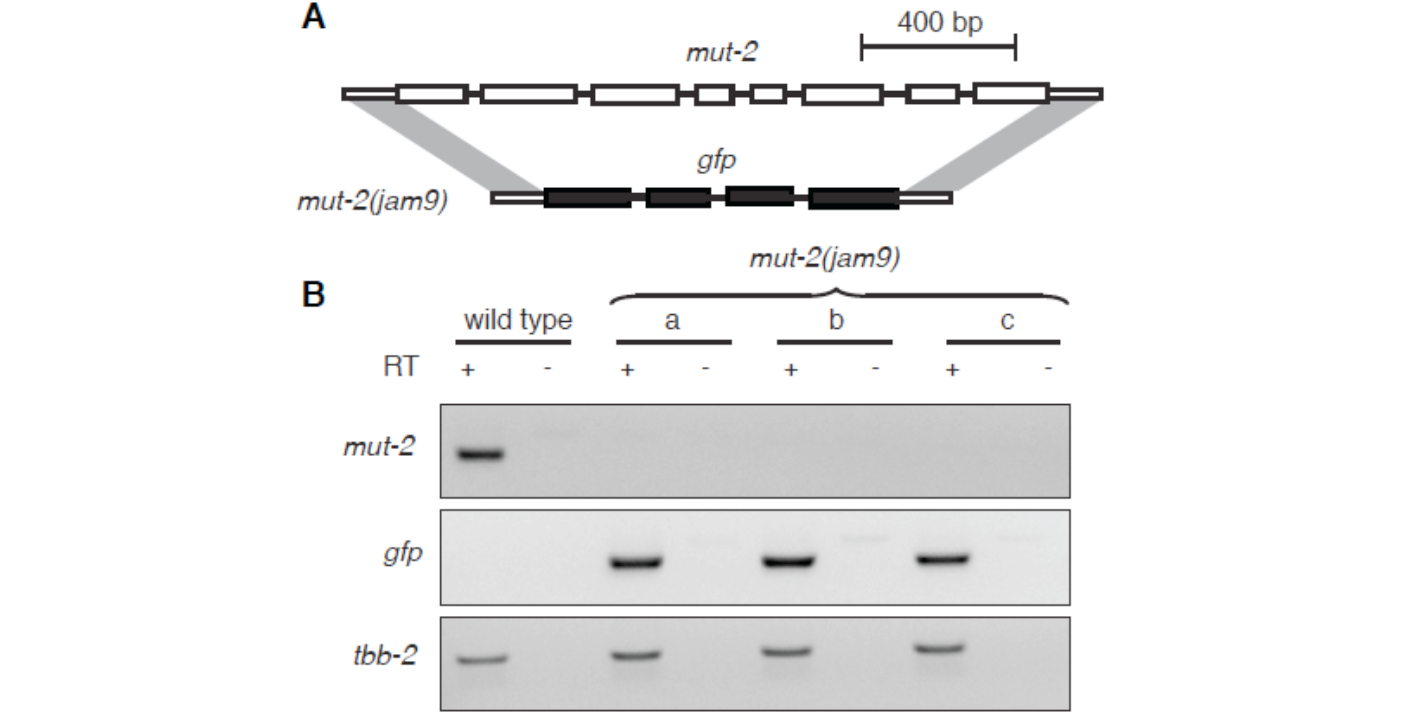
Generation of a *mut-2* null mutant using genome editing. (**A**) Schematic showing replacement of all exons (white boxes) and introns (lines) of *mut-2* with exons (black boxes) and introns (lines) of *gfp* using flanking regions of homology (grey shading) to generate a null mutant (*mut-2(jam9)*). **(B)** Transcript for *mut-2* is not detectable in three independent passages of *mut-2(jam9)*. PCR was used to amplify portions of *mut-2*, *gfp,* or *tbb-2* (control) in the presence (+) or absence (-) of reverse transcription (RT) of total RNA prepared from wild-type animals and from three independent passages (a, b, and c) of *mut-2(jam9)* animals.

**Supplemental Figure 3.**
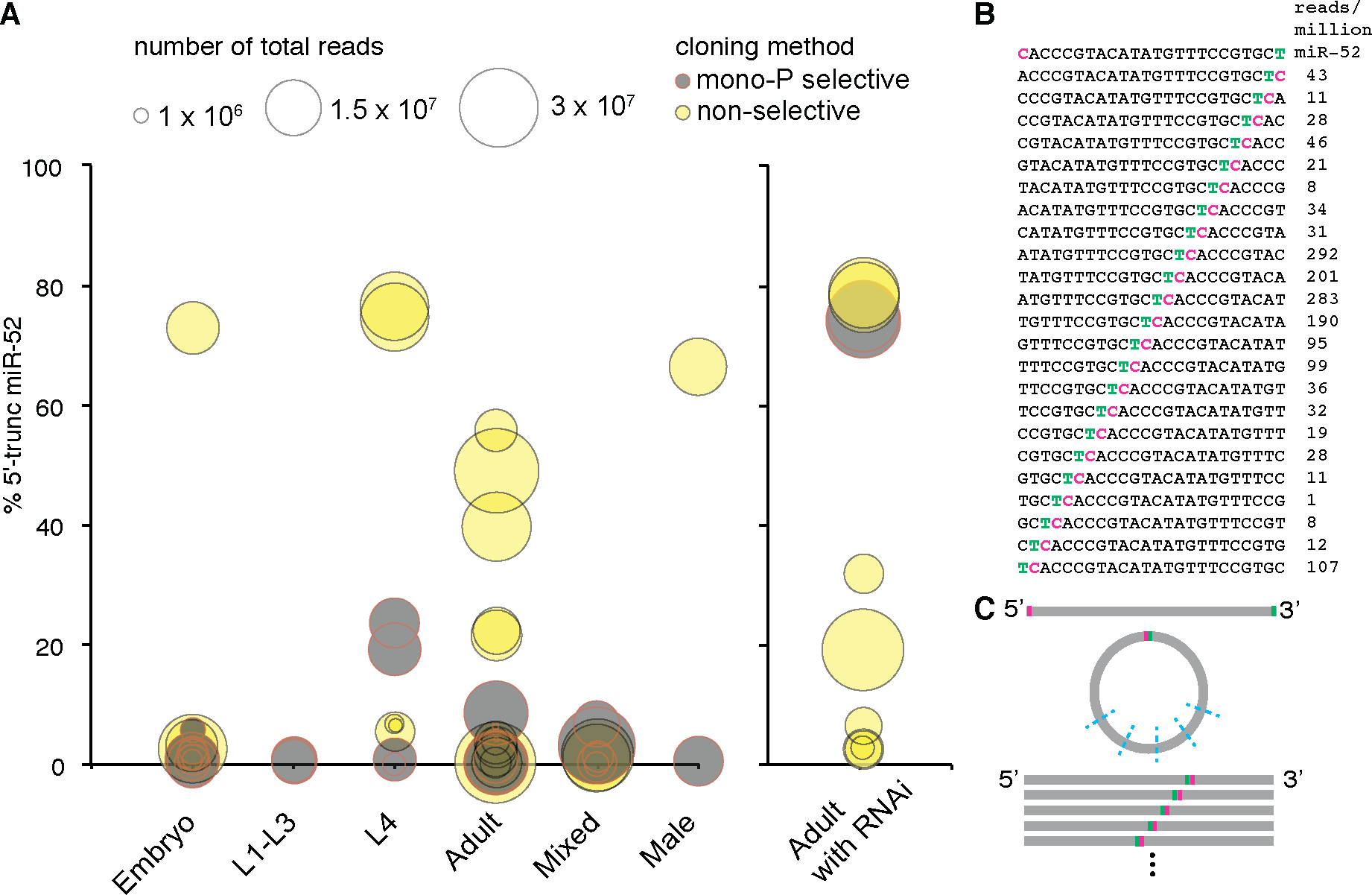
5’-truncated miR-52 was detected in many wild-type RNA-seq datasets.(**A**) Multiple wild-type datasets show shorter versions of miR-52 missing 5’ end nts. Small RNA-seq datasets from multiple labs prepared from wild-type worms (embryo, L1-L3, L4, Adult, mixed, male, or adult with RNAi) using mono-P selective (grey) or non-selective (yellow) cloning methods were examined for 5’ end variation in miR-52 length. The proportion of miR-52 (accession number MIMAT0000023) reads with complete 3’ end sequences that were missing any number of 5’ end nts (% 5’-truncated miR-52) were determined and plotted for each dataset. The size of the circle represents the total number of reads in the RNA-seq dataset. Also see Supplemental Table 1. (**B**) Circularly permuted version of miR-52 can be detected in RNA-seq datasets. All possible circularly permuted variants of miR-52 (top line) were detected in a wild-type dataset. The termini of the genomic miR-52 sequence are indicated in color (5’, magenta; 3’, green). Also see materials and methods. (**C**) Model for the generation of circularly permuted miRNA reads. Circularization and subsequent linearization (blue dashed lines) prior to adaptor ligation could explain the presence of circularly permuted miRNA reads in RNA-seq datasets.

**Supplemental Figure 4.**
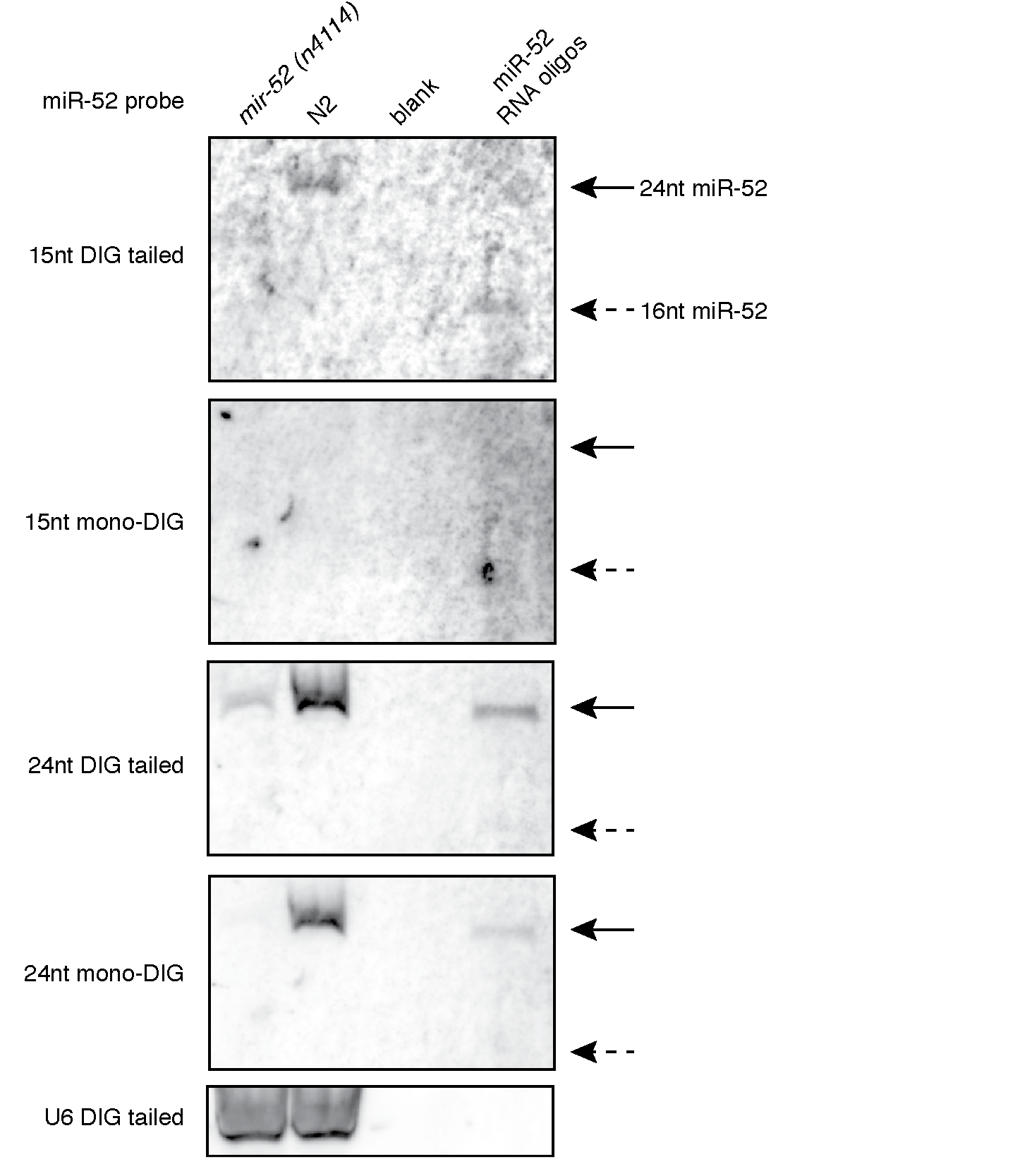
Comparison of mono-DIG and DIG-tailed probes hybridized at room temperature. Total RNA from wild type and miR-52 deletion animals was probed with 3’-mono-DIG and 3’-DIG tailed short and full-length probes hybridized at room temperature. Synthetic RNA representing 24 nt (10 fmol, solid arrow) and 16 nt (50 fmol, dashed arrow) of miR-52 sequence was used as size markers. U6 was probed to indicate loading.

**Supplemental Table 1.** RNA-seq datasets analyzed.

| Datasets used for Figure 1 |  |  |  |  |
| --- | --- | --- | --- | --- |
| SRX# | Corresponding Author | Allele | PMID |  |
| SRX013291 | Darryl Conte and Craig Mello | *mut-2(ne3364)* | 19800275 |  |
| SRX530098 | Gary Ruvkun and Taiowa Montgomery | *mut-2(ne298)* | 24684932 |  |
| SRR087429 | Gary Ruvkun | *mut-16(pk710)* | 21245313 |  |
| SRX136700 | Gary Ruvkun | N2 P-ind RNAi | 22542102 |  |
| SRX136689 | Gary Ruvkun | N2 P-ind | 22542102 |  |
| Datasets used for Supplemental Figure 3A |  |  |  |  |
| SRX# | Corresponding Author | Cloning | PMID | Minimum read count of 5'-trunc miR-52 for inclusion in Figure 3A |
| SRX136703 | Gary Ruvkun | P-dep RNAi | 22542102 | 40 |
| SRX136704 | Gary Ruvkun | P-dep RNAi | 22542102 | 40 |
| SRX129666 | Ho-Yi Mak | P-dep | 22508728 | 20 |
| SRX129660 | Ho-Yi Mak | P-dep | 22508728 | 20 |
| SRX424505 | Greg Hannon | P-dep | 24696458 | 21 |
| SAMN02195957 | Frank Slack | P-dep | 21129974 | 2 |
| SRX012407 | Nicholas Rajewsky | P-dep | 19734907 | 5 |
| SAMN02195956 | Frank Slack | P-dep | 21129974 | 2 |
| SRX059216 | Brenda Bass | P-dep | 22673872 | 7 |
| SRX012403 | Nicholas Rajewsky | P-dep | 19734907 | 1 |
| SAMN00008661 | John Kim | P-dep | 19846761 | 4 |
| SAMN00008664 | John Kim | P-dep | 19846761 | 2 |
| SRX012406 | Nicholas Rajewsky | P-dep | 19734907 | 4 |
| SRX012402 | Nicholas Rajewsky | P-dep | 19734907 | 1 |
| SRX003790 | Frank Slack | P-dep | 19460142 | 9 |
| SRX003786 | Frank Slack | P-dep | 19460142 | 10 |
| SRX003788 | Frank Slack | P-dep | 19460142 | 21 |
| SRX003792 | Frank Slack | P-dep | 19460142 | 21 |
| SRX003787 | Frank Slack | P-dep | 19460142 | 22 |
| SRX003789 | Frank Slack | P-dep | 19460142 | 21 |
| SRX003791 | Frank Slack | P-dep | 19460142 | 9 |
| SRX012405 | Nicholas Rajewsky | P-dep | 19734907 | 3 |
| SRX033023 | Frank Slack | P-dep |  | 5 |
| SRX012404 | Nicholas Rajewsky | P-dep | 19734907 | 34 |
| SRX695149 | Arnab Mukhopadhyay | P-dep | 25504288 | 5 |
| SRX059215 | Brenda Bass | P-dep | 22673872 | 7 |
| SAMN02196542 | Andrew Fire | P-dep | 20116306 | 32 |
| SRX019243 | Rene Ketting | P-dep | 20386745 | 6 |
| SRX095334 | Rene Ketting | P-dep | 22829772 | 10 |
| SRX192986 | Christoph Dieterich | P-dep | 23729632 | 10 |
| SRX136699 | Gary Ruvkun | P-ind RNAi | 22542102 | 20 |
| SRX136700 | Gary Ruvkun | P-ind RNAi | 22542102 | 20 |
| SRX113626 | Andrew Fire | P-ind RNAi | 22231482 | 20 |
| SRX113612 | Andrew Fire | P-ind RNAi | 22231482 | 15 |
| SRX113624 | Andrew Fire | P-ind RNAi | 22231482 | 10 |
| SRX113623 | Andrew Fire | P-ind RNAi | 22231482 | 6 |
| SRX113625 | Andrew Fire | P-ind RNAi | 22231482 | 22 |
| SRX113622 | Andrew Fire | P-ind RNAi | 22231482 | 10 |
| SRX111400 | Gary Ruvkun | P-ind | 22536158 | 20 |
| SRX111399 | Gary Ruvkun | P-ind | 22536158 | 10 |
| SRX193362 | Gary Ruvkun | P-ind | 23363624 | 22 |
| SRX193363 | Gary Ruvkun | P-ind | 23363624 | 20 |
| SRX446476 | Gary Ruvkun | P-ind | 24684932 | 21 |
| SRX193361 | Gary Ruvkun | P-ind | 23363624 | 20 |
| SRX446474 | Gary Ruvkun | P-ind | 24684932 | 20 |
| SRX129663 | Ho-Yi Mak | P-ind | 22508728 | 31 |
| SRX129669 | Ho-Yi Mak | P-ind | 22508728 | 20 |
| SRX142904 | David Bartel | P-ind | 22707570 | 10 |
| SAMN02196497 | Andrew Fire | P-ind | 20116306 | 1 |
| SAMN02196499 | Andrew Fire | P-ind | 20116306 | 5 |
| SRX328188 | Eric Miska | P-ind | 24696457 | 2 |
| SRX328190 | Eric Miska | P-ind | 24696457 | 5 |
| SRX136693 | Gary Ruvkun | P-ind | 22542102 | 10 |
| SRX136689 | Gary Ruvkun | P-ind | 22542102 | 21 |
| SRX098616 | Gary Ruvkun | P-ind | 22102828 | 20 |
| SRX098615 | Gary Ruvkun | P-ind | 22102828 | 20 |
| SRX189088 | Eric Miska | P-ind | 23811144 | 21 |
| SRX189089 | Darryl Conte and Craig Mello | P-ind | 23811144 | 22 |
| SRX017047 | Craig Mello & Darryl Conte Jr. | P-ind | 20133583 | 21 |
| SRX154615 | Craig Mello | P-ind | 22738724 | 21 |
| SRX181333 | Eric Miska | P-ind | 25258416 | 21 |
| Dataset used for Supplemental Figure 3B |  |  |  |  |
| **SRX#** | **Corresponding Author** | **PMID** |  |  |
| SRX136693 | Gary Ruvkun | 22542102 |  |  |

**Supplemental Table 2.**
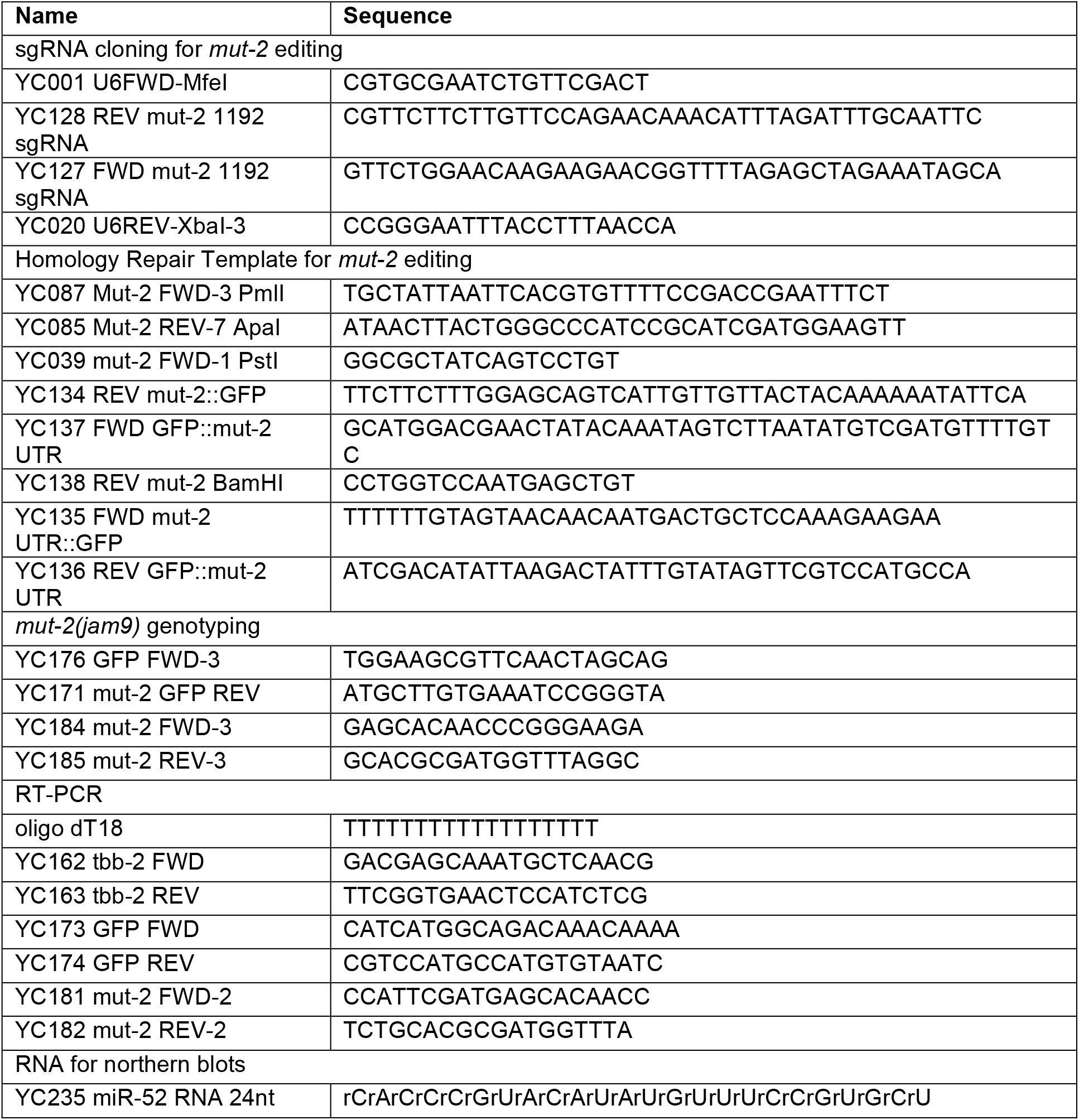
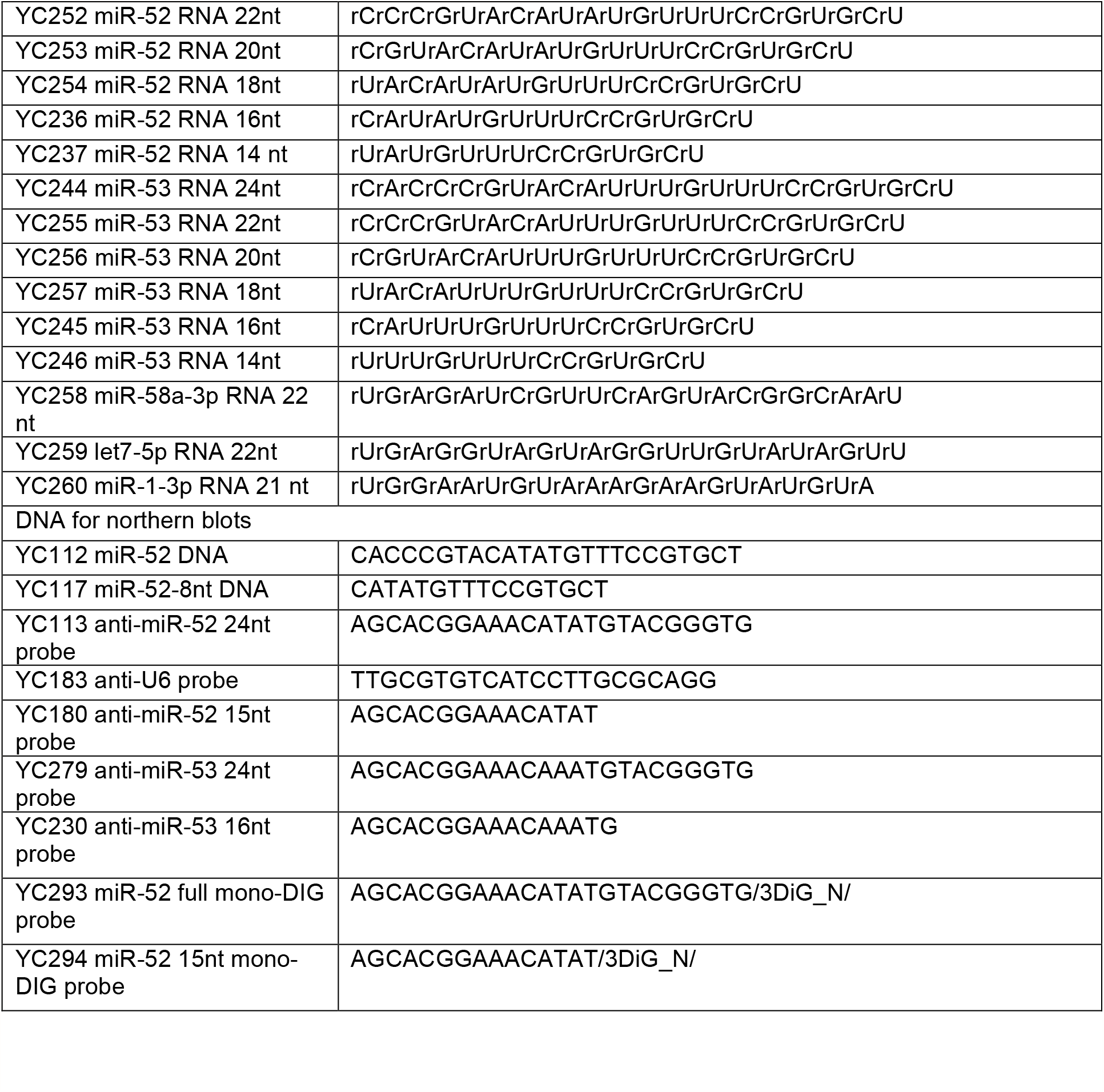
DNA and RNA oligos used.

## REFERENCES

1. Hammond, S.M. (2015) An overview of microRNAs. Adv. Drug Deliv. Rev. 87, 3–14.

2. Ozsolak, F. and Milos, P. (2011) RNA sequencing: advances, challenges and opportunities. Nat. Rev. Genetics, 12, 87–98.

3. Li, Y. and Kowdley, K. (2012) MicroRNAs in Common Human Diseases. Genomics Proteomics Bioinformatics, 10, 246–53.

4. Williams, Z., Ben-Dov, I., Elias, R., Mihailovic, A., Brown, M., Rosenwaks, Z. and Tuschl, T. (2013) Comprehensive profiling of circulating microRNA via small RNA sequencing of cDNA libraries reveals biomarker potential and limitations. Proc. Natl. Acad. Sci. U.S.A., 110, 4255–60.

5. Cheng, L., Sharples, R., Scicluna, B. and Hill, A. (2014) Exosomes provide a protective and enriched source of miRNA for biomarker profiling compared to intracellular and cell-free blood. J .Extracell. Vesicles, 3, 23743.

6. Pritchard, C.C., Cheng, H.H., Tewari, M. (2012) MicroRNA profiling: approaches and considerations. Nat. Rev. Genet. 13, 358–69.

7. Landgraf, P., Rusu, M., Sheridan, R., Sewer, A., Iovino, N., Aravin, A., Pfeffer, S., Rice, A., Kamphorst, A., Landthaler, M., et al. (2007) A mammalian microRNA expression atlas based on small RNA library sequencing. Cell, 129, 1401–14.

8. Morin, R., O’Connor, M., Griffith, M., Kuchenbauer, F., Delaney, A., Prabhu, A., Zhao, Y., McDonald, H., Zeng, T., Hirst, M. et al. (2008) Application of massively parallel sequencing to microRNA profiling and discovery in human embryonic stem cells. Genome Res., 18, 610–621.

9. Lagos-Quintana, M., Rauhut, R., Lendeckel, W., and Tuschl, T. (2001) Identification of novel genes coding for small expressed RNAs. Science. 294, 853–8.

10. Lau, N.C., Lim, L.P., Weinstein, E.G., and Bartel, D.P. (2001) An abundant class of tiny RNAs with probable regulatory roles in Caenorhabditis elegans. Science. 294, 858–62.

11. Lee, R.C., and Ambros, V. (2001) An extensive class of small RNAs in Caenorhabditis elegans. Science. 294, 862–4.

12. Neilsen, C.T., Goodall, G.J., and Bracken, C.P. (2012) IsomiRs — the overlooked repertoire in the dynamic microRNAome. Trends Genet. 28, 544–549.

13. Lee, M., Choi, Y., Kim, K., Jin, H., Lim, J., Nguyen, T.A., Yang, J., Jeong, M., Giraldez, A.J., Yang, H., Patel, D.J., and Kim, V.N. (2014) Adenylation of maternally inherited microRNAs by Wispy. Mol. Cell. 56, 696–707.

14. Katoh, T., Hojo, H., and Suzuki, T. (2015) Destabilization of microRNAs in human cells by 3’ deadenylation mediated by PARN and CUGBP Nucleic Acids Res., 43, 7521–34.

15. Marzi, M.J., Ghini, F., Cerruti, B., de Pretis, S., Bonetti, P., Giacomelli, C., Gorski, M.M., and Kress, T., Pelizzola, M., Muller, H., Amati, B., and Nicassio, F. (2016) Degradation dynamics of microRNAs revealed by a novel pulse-chase approach. Genome Res. 26, 554–65.

16. Lewis, B.P., Shih, I.H., Jones-Rhoades, M.W., Bartel, D.P., and Burge, C.B. (2003) Prediction of mammalian microRNA targets. Cell. 115, 787–98.

17. Fukunaga, R., Han, B., Hung, J., Xu, J., Weng, Z. and Zamore, P. (2012) Dicer partner proteins tune the length of mature miRNAs in flies and mammals. Cell, 151, 533–46.

18. Heo, I., Ha, M., Lim, J., Yoon, M., Park, J., Kwon, S., Chang, H. and Kim VN. (2012) Mono-uridylation of pre-microRNA as a key step in the biogenesis of group II let-7 microRNAs. Cell, 151, 521–32.

19. Lee, H. and Doudna, J.A. (2012) TRBP alters human precursor microRNA processing in vitro. RNA, 18, 2012–2019.

20. Tan, G., Chan, E., Molnar, A., Sarkar, R., Alexieva, D., Isa, I., Robinson, S., Zhang, S., Ellis, P., Langford, C., et al. (2014) 5’ isomiR variation is of functional and evolutionary importance. Nucleic Acids Res., 42, 9424–9435.

21. Raabe, C.A., Tang,T.-H., Brosius, J., Rozhdestvensky, T.S. (2014) Biases in small RNA deep sequencing data. Nucleic Acids Res., 42, 1414–1426.

22. Dabney, J. and Meyer, M. (2012) Length and GC-biases during sequencing library amplification: a comparison of various polymerase-buffer systems with ancient and modern DNA sequencing libraries. Biotechniques, 52, 87–94.

23. Orpana, A.K., Ho, T.H. and Stenman, J. (2012) Multiple heat pulses during PCR extension enabling amplification of GC-rich sequences and reducing amplification bias. Anal. Chem., 84, 2081–2087.

24. Hafner, M., Renwick, N., Brown, M., Mihailovi?, A., Holoch, D., Lin, C., Pena, J., Nusbaum, J., Morozov, P., Ludwig, J., et al., (2011) RNA-ligase-dependent biases in miRNA representation in deep-sequenced small RNA cDNA libraries. RNA, 17, 1697–712.

25. Harris, C., Molnar, A., Müller, S. and Baulcombe, D. (2015) FDF-PAGE: a powerful technique revealing previously undetected small RNAs sequestered by complementary transcripts. Nucleic Acids Res., 43, 7590–7599.

26. Kim, Y., Yeo, J., Kim, B., Ha, M. and Kim, V.N. (2012) Short structured RNAs with low GC content are selectively lost during extraction from a small number of cells. Mol. Cell, 46, 893–5.

27. Válóczi, A., Hornyik, C., Varga, N., Burgyán, J., Kauppinen, S., and Havelda, Z. (2004) Sensitive and specific detection of microRNAs by northern blot analysis using LNA-modified oligonucleotide probes. Nucleic Acids Res. 32, e175.

28. Pall, G.S., Codony-Servat, C., Byrne, J., Ritchie, L. and Hamilton, A. (2007) Carbodiimide-mediated cross-linking of RNA to nylon membranes improves the detection of siRNA, miRNA and piRNA by northern blot. Nucleic Acids Res., 35, e60.

29. Höltke, H.J. and Kessler, C. (1990) Non-radioactive labeling of RNA transcripts in vitro with the hapten digoxigenin (DIG); hybridization and ELISA-based detection. Nucleic Acids Res. 18, 5843–51.

30. Ramkissoon, S.H., Mainwaring, L.A., Sloand, E.M., Young, N.S., and Kajigaya, S. (2006) Nonisotopic detection of microRNA using digoxigenin labeled RNA probes. Mol Cell Probes. 20, 1–4.

31. Kim, S.W., Li, Z., Moore, P.S., Monaghan, A.P., Chang, Y., Nichols, M., and John, B. (2010) A sensitive non-radioactive northern blot method to detect small RNAs. Nucleic Acids Res., 38, e98.

32. Schwarzkopf, M., and Pierce, N.A. (2016) Multiplexed miRNA northern blots via hybridization chain reaction. Nucleic Acids Res. e-pub Jun 7.

33. Giardine, B., Riemer, C., Hardison, R.C., Burhans, R., Elnitski, L., Shah, P., Zhang, Y., Blankenberg, D., Albert, I., Taylor, J., et al., (2005) Galaxy: a platform for interactive large-scale genome analysis. Genome Res., 15, 1451–5.

34. Blankenberg, D., Von Kuster, G., Coraor, N., Ananda, G., Lazarus, R., Mangan, M., Nekrutenko, A., and Taylor, J. (2010) Galaxy: a web-based genome analysis tool for experimentalists. Curr. Prot. Mol. Biol., 19, 1–21.

35. Goecks, J., Nekrutenko, A., Taylor, J., and The Galaxy Team. (2010) Galaxy: a comprehensive approach for supporting accessible, reproducible, and transparent computational research in the life sciences. Genome Biol., 11, R86.

36. Blankenberg, D., Von Kuster, G., Bouvier, E., Baker, D., Afgan, E., Stoler, N., Galaxy Team, Taylor, J., and Nekrutenko, A. (2014) Dissemination of scientific software with Galaxy ToolShed. Genome Biol. 15, 403.

37. Chen, C., Simard, M., Tabara, H., Brownell, D., McCollough, J. and Mello, C. (2005) A member of the polymerase beta nucleotidyltransferase superfamily is required for RNA interference in C. elegans. Curr. Biol., 15, 378–83.

38. Dickinson, D., Ward, J., Reiner, D. and Goldstein, B. (2013) Engineering the Caenorhabditis elegans genome using Cas9-triggered homologous recombination. Nat. Methods, 10, 1028–34.

39. Friedland, A., Tzur, Y., Esvelt, K., Colaiácovo, M., Church, G. and Calarco, J. (2013) A Heritable genome editing in C. elegans via a CRISPR-Cas9 system. Nat. Methods, 10, 741–3.

40. Heigwer, F., Kerr, G. and Boutros, M. (2014) E-CRISP: fast CRISPR target site identification. Nat.Methods, 11, 122–123.

41. Frøkjaer-Jensen, C., Davis, M., Hopkins, C., Newman, B., Thummel, J., Olesen, S., Grunnet, M. and Jorgensen, E. (2008) Single-copy insertion of transgenes in Caenorhabditis elegans. Nat. Genetics, 40, 1375–83

42. Voutev, R. and Hubbard, E. (2008) A “FLP-Out” system for controlled gene expression in Caenorhabditis elegans. Genetics, 180, 103–119.

43. Frøkjær-Jensen, C., Davis, M., Ailion, M. and Jorgensen, E. (2012) Improved Mos1-mediated transgenesis in C. elegans. Nat. Methods, 9, 117–8.

44. Schneider, C.A., Rasband, W.S. and Eliceiri, K.W. (2012) NIH Image to ImageJ: 25 years of image analysis. Nat. Methods 9, 671–5.

45. Brenner, S. (1974) The genetics of Caenorhabditis elegans. Genetics, 77, 71–94.

46. Zhang, C., Montgomery, T., Gabel, H., Fischer, S., Phillips, C., Fahlgren, N., Sullivan, C., Carrington, J. and Ruvkun, G. (2011) mut-16 and other mutator class genes modulate 22G and 26G siRNA pathways in Caenorhabditis elegans. Proc. Natl. Acad. Sci. U.S.A., 108, 1201–1208.

47. Phillips, C., Montgomery, T., Breen, P. and Ruvkun, G. (2012) MUT-16 promotes formation of perinuclear mutator foci required for RNA silencing in the C. elegans germline. Genes Dev., 26, 1433–44.

48. Alvarez-Saavedra, E. and Horvitz, H.R. (2010) Many families of C. elegans microRNAs are not essential for development or viability. Curr. Biol. 20, 367–73.

49. Batista, P., Ruby, J., Claycomb, J., Chiang, R., Fahlgren, N., Kasschau, K., Chaves, D., Gu, W., Vasale, J., Duan, S., et al. (2008) PRG-1 and 21U-RNAs interact to form the piRNA complex required for fertility in C. elegans. Mol. Cell, 31, 67–78.

50. Collins, J., Saari, B. and Anderson, P. (1987) Activation of a transposable element in the germ line but not the soma of Caenorhabditis elegans. Nature, 328, 726–728.

51. Beckmann, B.M., Grünweller, A., Weber, M.H. and Hartmann, R.K. (2010) Northern blot detection of endogenous small RNAs (approximately14 nt) in bacterial total RNA extracts. Nucleic Acids Res. 38, e147.

52. Shaw, W., Armisen, J., Lehrbach, N. and Miska, E. (2010) The Conserved miR-51 microRNA Family Is Redundantly Required for Embryonic Development and Pharynx Attachment in Caenorhabditis elegans. Genetics, 185, 897–905.

53. Chatterjee, S. and Großhans, H. (2009) Active turnover modulates mature microRNA activity in Caenorhabditis elegans. Nature. 461, 546–9.

54. Miki, T.S., Rüegger, S., Gaidatzis, D., Stadler, M.B. and Großhans, H. (2014) Engineering of a conditional allele reveals multiple roles of XRN2 in Caenorhabditis elegans development and substrate specificity in microRNA turnover. Nucleic Acids Res. 42, 4056–67.

55. Memczak, S., Jens, M., Elefsinioti, A., Torti, F., Krueger, J., Rybak, A., Maier, L., Mackowiak, S., Gregersen, L., Munschauer, M. et al. (2013) Circular RNAs are a large class of animal RNAs with regulatory potency. Nature, 495, 333–338.

56. Kosmaczewski, S., Edwards, T., Han, S., Eckwahl, M., Meyer, B., Peach, S., Hesselberth, J., Wolin, S. and Hammarlund, M. (2014) The RtcB RNA ligase is an essential component of the metazoan unfolded protein response. EMBO Rep., 15, 1278–85.

57. Wassarman, K.M. and Saecker, R.M. (2006) Synthesis-mediated release of a small RNA inhibitor of RNA polymerase. Science 314, 1601–1603.

58. Blumenfeld, A.L. and Jose, A.M. (2016) Reproducible features of small RNAs in C. elegans reveal NU RNAs and provide insights into 22G RNAs and 26G RNAs. RNA 22, 184–92.

59. Li, Z., Kim, S.W., Lin, Y., Moore, P.S., Chang, Y., and John, B. (2009) Characterization of viral and human RNAs smaller than canonical MicroRNAs. J. Virol. 83, 12751–12758.

60. Wang, X. and Clark, A.G. (2014) Using next-generation RNA sequencing to identify imprinted genes. Heredity (Edinb). 113, 156–166.

61. Strerath, M., Detmer, I., Gaster, J., and Marx, A. (2007) Modified oligonucleotides as tools for allele-specific amplification. Methods Mol. Biol. 402, 317–28.

62. Rüegger, S. and Großhans, H. (2012) MicroRNA turnover: when, how, and why. Trends Biochem. Sci. 37, 436–446.

